# State of the Interactomes: an evaluation of molecular networks for generating biological insights

**DOI:** 10.1101/2024.04.26.587073

**Authors:** Sarah N. Wright, Scott Colton, Leah V. Schaffer, Rudolf T. Pillich, Christopher Churas, Dexter Pratt, Trey Ideker

## Abstract

Advancements in genomic and proteomic technologies have powered the use of gene and protein networks (“interactomes”) for understanding genotype-phenotype translation. However, the proliferation of interactomes complicates the selection of networks for specific applications. Here, we present a comprehensive evaluation of 46 current human interactomes, encompassing protein-protein interactions as well as gene regulatory, signaling, colocalization, and genetic interaction networks. Our analysis shows that large composite networks such as HumanNet, STRING, and FunCoup are most effective for identifying disease genes, while smaller networks such as DIP and SIGNOR demonstrate strong interaction prediction performance. These findings provide a benchmark for interactomes across diverse network biology applications and clarify factors that influence network performance. Furthermore, our evaluation pipeline paves the way for continued assessment of emerging and updated interaction networks in the future.

## INTRODUCTION

Molecular networks (“interactomes”) are a crucial tool in biomedical research for translating complex biological data into actionable insights. These networks are constructed from diverse genomic, proteomic, biochemical, and statistical data sources to represent known or measured biological interactions among genes, proteins, and other biological entities. Interactions encompass a wide range of types, including physical protein-protein interactions, regulatory relationships, signaling and metabolic pathways, and functional associations, each of which contribute to our understanding of molecular and cellular functions.

By leveraging the information in these interactomes, network biology provides insights into gene and protein functions at a systems level, thus enabling a comprehensive understanding of biological and disease processes. Such approaches have improved the interpretation of genome-wide association studies (GWAS)^1–4^, prioritized disease genes^5,6^, accelerated the discovery of gene functions^7,8^, and revealed functional similarities and differences between species and cell types^9,10^. The success of each approach can depend critically on selecting an appropriate interactome for study. However, the rapid generation of biological data has led to a proliferation of available networks, making network selection increasingly challenging.

No matter the type of analysis performed, the ability of a network approach to generate biological insights depends on what information is and is not present in an interactome. It is widely recognized that our knowledge of biological interactions is incomplete, especially for experimentally supported interactions^11–13^. However, the extent to which gaps are skewed towards certain genes or biological processes is less well understood. Many factors can contribute to this apparent skew, including experimental constraints or bias toward highly studied or highly expressed genes^14^. It is important to seek to understand these effects across interactomes, as any bias in an interactome is likely to be reflected in downstream analyses

Several years ago, we established methods for systematically evaluating human molecular networks and demonstrated that large composite interactomes provided the best performance for prioritizing disease genes^15^. This work culminated in the Parsimonious Composite Network (PCNet), a consensus network that includes the most supported interactions across different network resources while excluding potentially spurious relationships^15^. Given the continued increases in size and quantity of interactomes, ongoing benchmarking of molecular networks is essential for guiding network selection. Accordingly, here we provide an updated and expanded evaluation, covering the most extensive information to date on the contents and performance of human molecular networks currently available in the public domain. Concomitant with these efforts is the release of PCNet2.0, a new and improved consensus human gene network.

In addition to the established procedure for evaluating the performance of interactomes via disease gene prioritization, we introduce a second, interaction-centric, evaluation metric. This approach leverages interaction prediction algorithms, which provide an efficient method for addressing network incompleteness and have previously been used to identify and prioritize novel interactions^16,17^. Previously, benchmarking studies have assessed the performance of different interaction prediction algorithms across a small number of human interactomes^18,19^. Based on these studies, we now implement interaction prediction across a wide range of interactomes to assess the influence of the underlying network architecture on prediction accuracy. We define gold standard interaction sets for this evaluation and create a validation pipeline for novel predictions using AlphaFold-Multimer^20^.

Through this comprehensive evaluation, we aim to provide an up-to-date survey of 46 interactomes, equipping researchers with the information necessary to navigate network selection and thereby power the generation of biological insights across various applications. Our consensus networks are easily accessible via the Network Data Exchange (NDEx^21,22^) at www.ndexbio.org, and our evaluation pipeline is available at https://github.com/sarah-n-wright/Network_Evaluation_Tools, allowing for ongoing analysis of biological networks.

## RESULTS

### A plethora of interaction networks from diverse but overlapping sources

We performed a census of current biomolecular network resources, focusing on gene and protein-centric human interactomes. Our survey identified 46 publicly available networks (Supplemental Table 1), which we classified into three categories: Experimental - networks formed from a single experimental source, Curated - networks curated manually or computationally from literature sources, and Composite - networks directly incorporating multiple curated or experimental databases. Across these categories, we observed substantial diversity in the network features and data sources (Figure 1A). For example, while 93% of the interactomes incorporated physical protein-protein interactions (PPIs), less than 25% contained information from genome or protein structural similarities. The majority of network resources we surveyed (70%) contained interaction evidence from multiple species. We excluded all non-human interactions from these networks for our human-centric analysis, except where the authors explicitly used orthologous interactions to enrich human networks.

**Figure 1.**
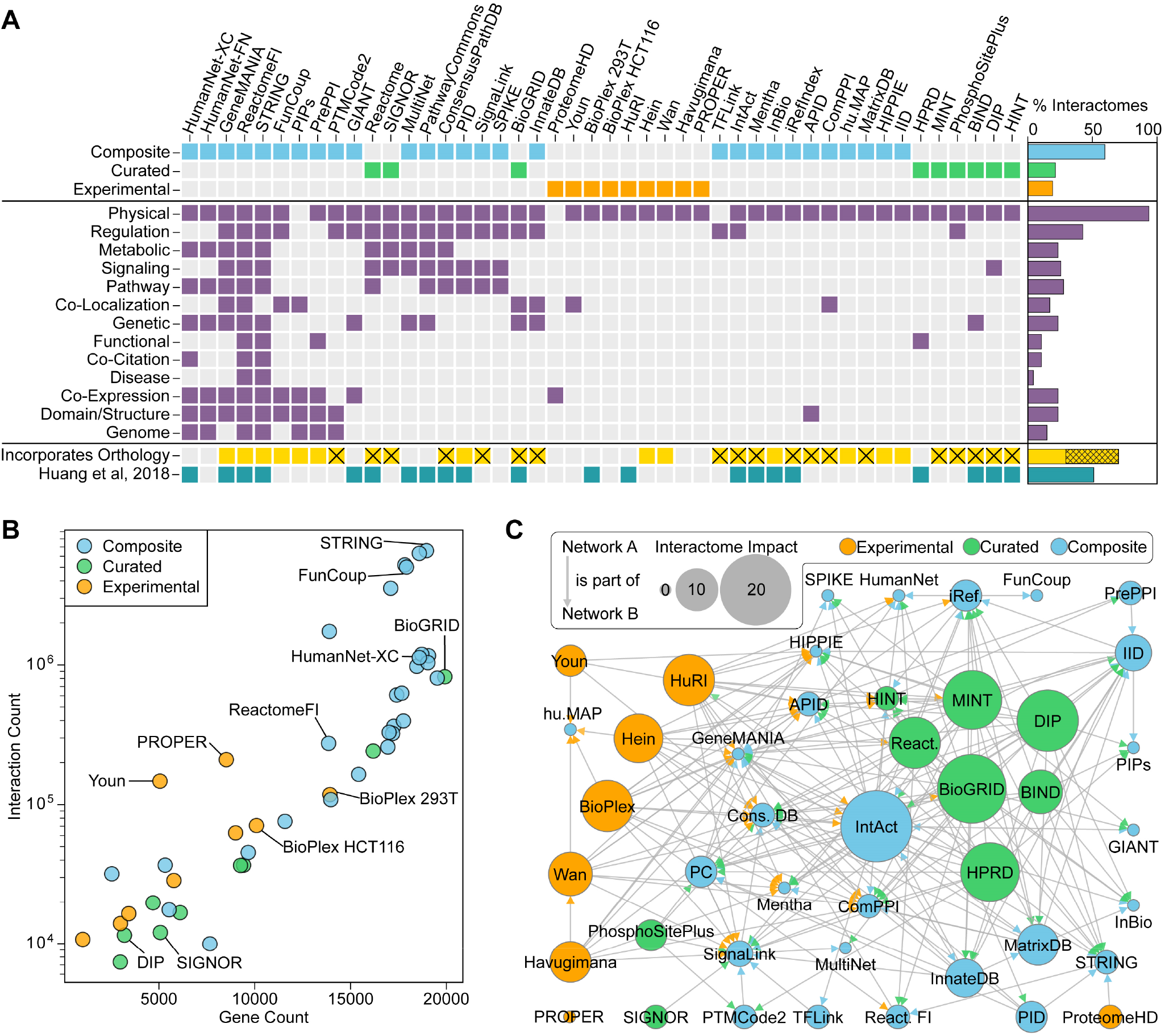
Network contents, sources, and dependencies. A) Interactome classification, interaction types, and network features, including whether a network was included in our previous evaluation (Huang et al, 2018). Crosses indicate that non-human interactions were filtered out in data processing. The bar chart shows the percentage of all interactomes with each feature. B) Interactome size plotted by the number of interactions versus the number of genes, with color indicating interactome classification. Gene and interaction counts represent unique values after data processing and conversion of identifiers to NCBI Gene IDs. C) Dependency relationships between interactomes, colored by classification. Arrows represent that the source interactome is incorporated into the target interactome. The arrow color indicates the classification of the source network. Node size indicates the “Interactome Impact,” defined as the number of times a given interactome is cited as a source by other evaluated network sources.

All networks were standardized through our improved network processing pipeline, achieving a reduction of over 60% in genes that could not be mapped to NCBI gene identifiers (Figure S1A). Composite networks tended to be much larger than experimental and curated networks, both in the number of genes represented and the number of reported interactions (Figure 1B). Building on the 21 networks analyzed in our prior publication, we updated 14 and extended the corpus with 25 additional interactomes^15^. Of the 14 updated networks, the Human Reference Interactome^23^ (HuRI) grew the most, with a 2.6-fold increase in genes and a 4.6-fold increase in interactions compared to the previously evaluated Human Interactome^24^ (HI-II-14) (Figure S1B). While the number of distinct databases has increased substantially, we found that most available networks rely on similar sources and have extensive interdependencies (Figure 1C). The network resources with the highest impact on other interactomes were BioGRID^25^ (19), IntAct^26^ (20), DIP^27^ (16), MINT^28^ (15), and HPRD^29–31^ (15).

### Gaps remain in coverage of the human proteome

Clearly, a network analysis can only discover genes and processes existing within the interactome selected for study. Despite over 99% of protein-coding genes (as defined by the HUGO Gene Nomenclature Committee, HGNC^32,33^) being represented in at least one interactome, we found that their distribution varies widely across networks (Figure 2A). Other genetic elements, such as non-coding RNAs and pseudogenes, are sparsely represented (Figure 2B). Looking at the relationship between gene presence in interactomes and gene citation frequency of protein-coding genes, we observed a significant correlation (Figure 2C, r_s,citation_ = 0.80, p < 1×10^−20^), which was diminished but not eliminated when considering only experimental networks (Figure S2A, r_s_ = 0.48, p < 1×10^−20^). High mRNA expression (GTEx) and protein abundance (Human Protein Atlas, HPA^34,35^) also significantly correlated with increased network coverage (r_s,mRNA_ = 0.63, r_s,protein_ = 0.40, p < 1×10^−20^, Figure 2D-E). After adjusting for mRNA and protein expression, the correlation between network coverage and gene citation was partly reduced (r_s,citation_ = 0.59, p < 1×10^−20^), indicating that expression levels contribute to, but do not completely explain, the observed citation bias (Figure S2B).

**Figure 2.**
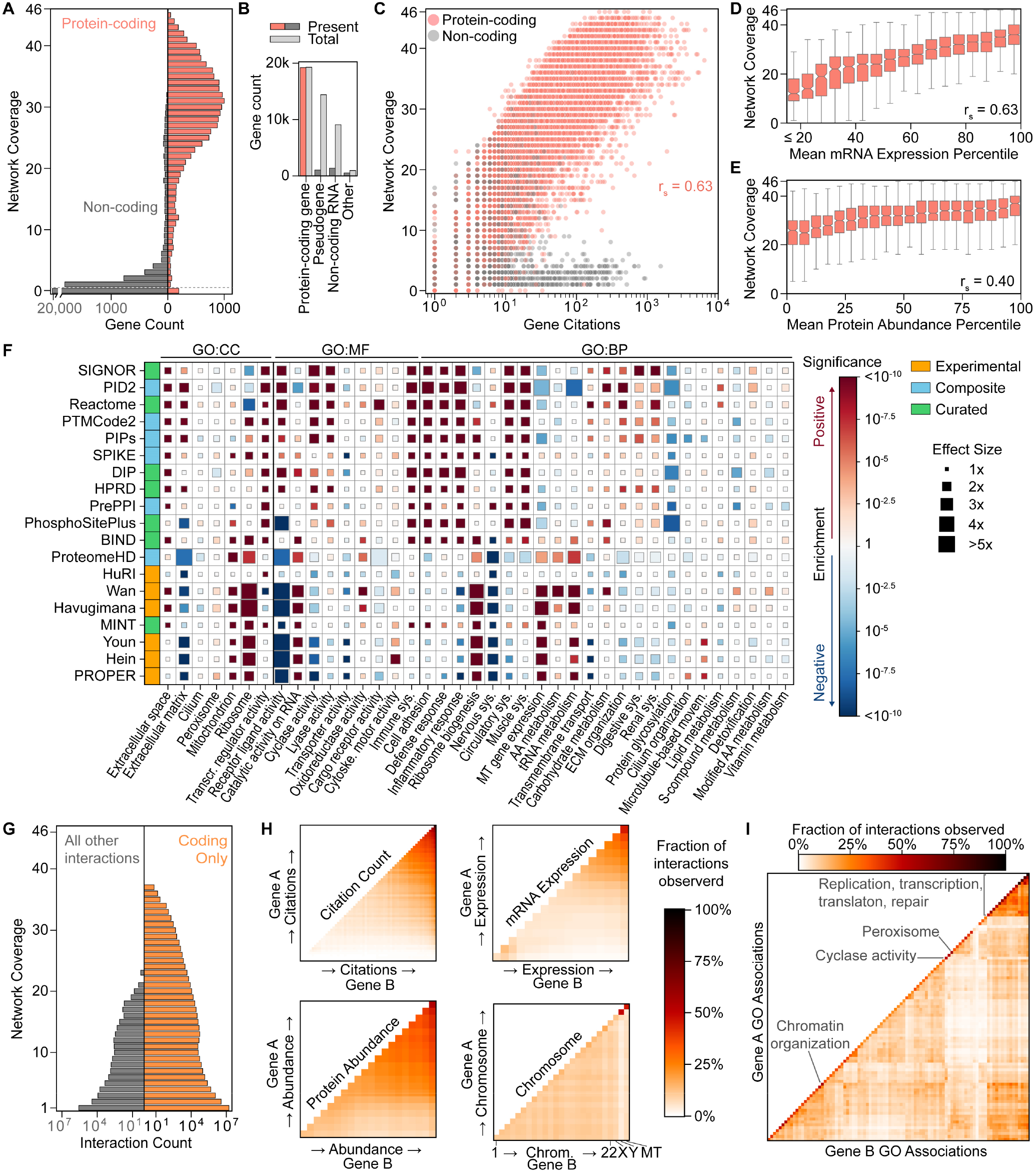
Representation analysis of interactome contents. A) Network coverage of protein-coding and non-coding genes, defined by HGNC locus type. B) Presence of genes in at least one interactome by HGNC locus type, compared to total genes of each type. C) Network coverage versus NCBI gene citation count (December 20, 2023). D-E) Network coverage of protein-coding genes as a function of mean mRNA expression across all tissues in GTEx and mean protein abundance across all tissues in the Human Protein Atlas (HPA). Box plots show the mean network coverage across expression percentiles. Outliers not shown. F) Functional enrichment of network genes across the Gene Ontology Slim. Networks with < 10,000 genes and terms with at least one significantly under-enriched network (q < 0.01) are shown. See also Supplemental Table 2. G) Network coverage of interactions between protein-coding genes (“Coding only”) and those involving at least one non-coding gene (“All other interactions”). H-I) Distribution of all interactions as a function of gene annotations. Genes are binned based on increasing citation count, mRNA expression percentile, protein abundance percentile, and chromosome (H) or GO Slim annotations (I). The fraction of observed interactions is calculated relative to the number of possible gene A and gene B combinations between bins.

Across all interactomes, 23.6 million unique interactions were reported, with 98% representing interactions between protein-coding genes. The majority of interactions (72.9%) were unique to a single network, while those reported in 14 or more networks comprised only 1% of the interactions (Figure 2G). High expression and citation levels were generally associated with high connectivity (Figure 2H), though increased connectivity was observed among a subset of genes with low mean mRNA expression - driven by interactions between genes with tissue-specific expression in whole blood, thyroid, and skin (Figure S2C). Connectivity was largely uniform within and between chromosomes, except for the Y and mitochondrial chromosomes (Figure 2H). Analysis of interaction frequency among Gene Ontology (GO) annotations highlighted extensive connectivity among genes involved in genomic organization and processing (Figure 2I). In contrast, genes associated with some functions and components, such as cyclase activity and the peroxisome, had more restricted interactions.

### Large composite networks remain the most powerful for disease gene prioritization

Next, we assessed each interactome’s ability to recover a range of disease-associated gene sets using a network propagation framework (Methods, Figure 3A). This approach is based on the principle that genes in close proximity in a biological network are likely to share biological functions. For disease gene set recovery, we used network propagation to rank all genes within an interactome based on their proximity to a known set of disease genes. We then assessed the position of held-out members of the gene set within these propagation ranks to determine how well the network structure recovered known disease associations. To quantify this gene set recovery performance, we compared these results to ranks generated with shuffled interactomes to generate performance Z-scores^15^. Our analysis evaluated gene set recovery performance for all interactomes using diverse gene sets sourced from DisGeNET^38^ (“literature”, n=904) and the GWAS catalog^39^ (“genetic”, n=439). Literature gene sets, being larger, generally showed better interactome coverage and gene set recovery performance (Figure S3A).

**Figure 3.**
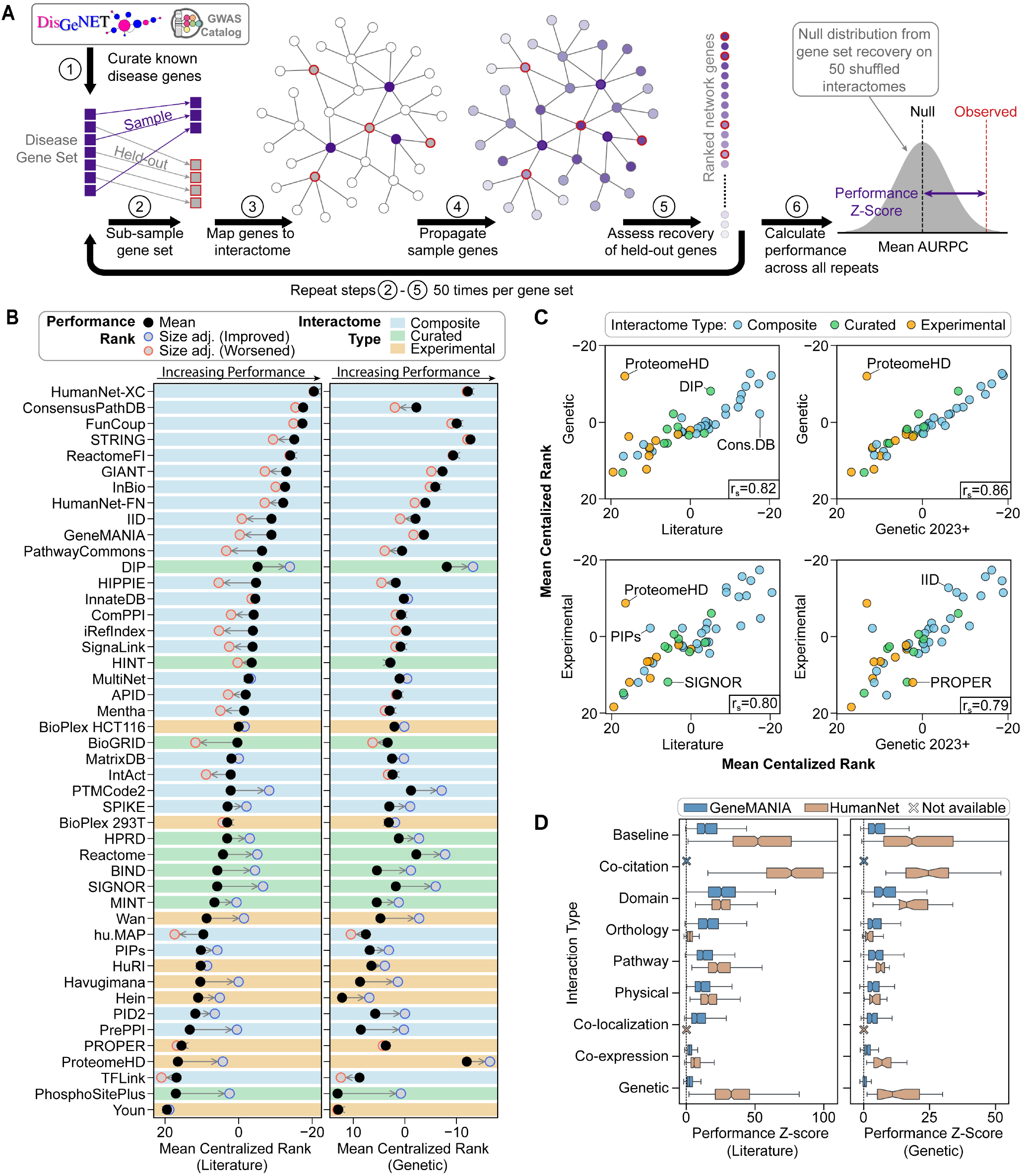
Evaluation of gene set recovery performance. A) Overview of the gene set recovery pipeline to generate performance Z-scores. B) Mean network performance centralized rank across all literature (left) and genetic (right) gene sets. A more negative rank indicates better relative performance. Arrows and colored circles show the mean size-adjusted rank, and background indicates network classification. C) Spearman correlations of interactome centralized ranks between literature, genetic, genetic 2023+, and experimental gene sets. Genetic 2023+ gene sets were defined from GWAS published July 27, 2023, or later. Experimental gene sets were defined from publications dated after January 1, 2024. D) Gene set recovery performance of interaction type-specific networks from HumanNet and GeneMANIA. See also Supplemental Table 3.

Ranking the performance Z-scores across all gene sets revealed that large composite networks HumanNet-XC^40^ and STRING^41^ were most effective for prioritizing literature and genetic disease genes, respectively (Figure 3B). Given the previously established relationship between network size and gene set recovery performance^15^, we also computed size-adjusted performance metrics by regressing the number of interactions per network. Smaller networks such as DIP^27^, SIGNOR^36^, and PhosphositePlus^42^, rose in the rankings after adjustment, indicating that their interactions are highly informative on a per-interaction basis (Figure 3B).

Interactome performance was consistent across literature and genetic gene sets (r_s_ = 0.82, p = 4.7×10^−12^), with exceptions for ConsensusPathDB^43^, DIP^27^, and ProteomeHD^44^, which showed large discrepancies across gene set sources (Figure 3C). These exceptions indicate that some interactomes are better suited to analyzing certain data types. To address potential circularity in our analysis due to literature curation - a concern that arises because some gene sets and interactomes may be derived from overlapping literature sources - we also assessed the performance of additional gene sets derived from GWAS published after July 2023 (“genetic 2023+”, n=48) and experimental studies published after January 1, 2024 (“experimental”, n=17). These gene sets were thus generated from data available only after the release of any evaluated interactome that utilized text mining of biomedical literature. Performance rankings for genetic gene sets were highly correlated with those for genetic 2023+ gene sets (r_s_ = 0.86, p = 3.1×10^−14^, Figure 3C), indicating minimal bias from literature curation for genetic gene sets. Correspondence to experimental gene sets was also positively correlated, though some interactomes, such as ConsensusPathDB^43^, showed reduced performance for non-literature gene sets. Overall, the strong correlations in network rankings across different gene sets underscored the robustness of our analysis against concerns of circularity due to literature curation.

### Not all interactions are created equal

Given that the highest-performing networks were composite networks comprising many evidence types, we examined the contributions of the various interaction types in two large network databases: HumanNet v3^40^ and GeneMANIA^45^. We defined interaction-type specific networks from each source, ranging from the small HumanNet-PG (phylogenetic) network to the large GeneMANIA co-expression network (Figure S3B). The two physical interaction networks were highly similar (Jaccard = 0.58), representing a high degree of consistency in physical interaction definitions, especially compared to other interaction types (Figure S3C). Analysis of gene set recovery performance revealed that domain similarity, pathway, and physical interactions were particularly informative across both HumanNet and GeneMANIA (Figure 3D). In contrast, results for genetic, orthologous, and co-expression interactions were inconsistent between the two sources, reflecting the influence of differing network construction procedures. The HumanNet co-citation network outperformed the full HumanNet-XC interactome in recovering literature gene sets, likely indicating some circularity between the literature-focused network and literature-derived gene sets.

### Consensus networks enhance gene set recovery performance

As part of our earlier systematic evaluation of molecular networks by Huang et al.^15^, we demonstrated that creating consensus networks, such as the Parsimonious Consensus Network (PCNet 1.0), could enhance gene set recovery performance. To assess the ongoing benefits of combining interactomes, we evaluated two approaches to assembling consensus networks: “global” composites and “ranked” composites (Figure 4A). Global composite networks were constructed following the methodology of PCNet 1.0, in which a progressive series of composites was constructed by requiring interaction coverage from an increasing number of the 46 interactomes evaluated. Alternatively, we constructed ranked composite networks by varying the number of top-ranked interactomes considered, based on median size-adjusted performance. For each ranked composite network threshold, we created two networks requiring interaction coverage from at least two or three interactomes, respectively.

**Figure 4.**
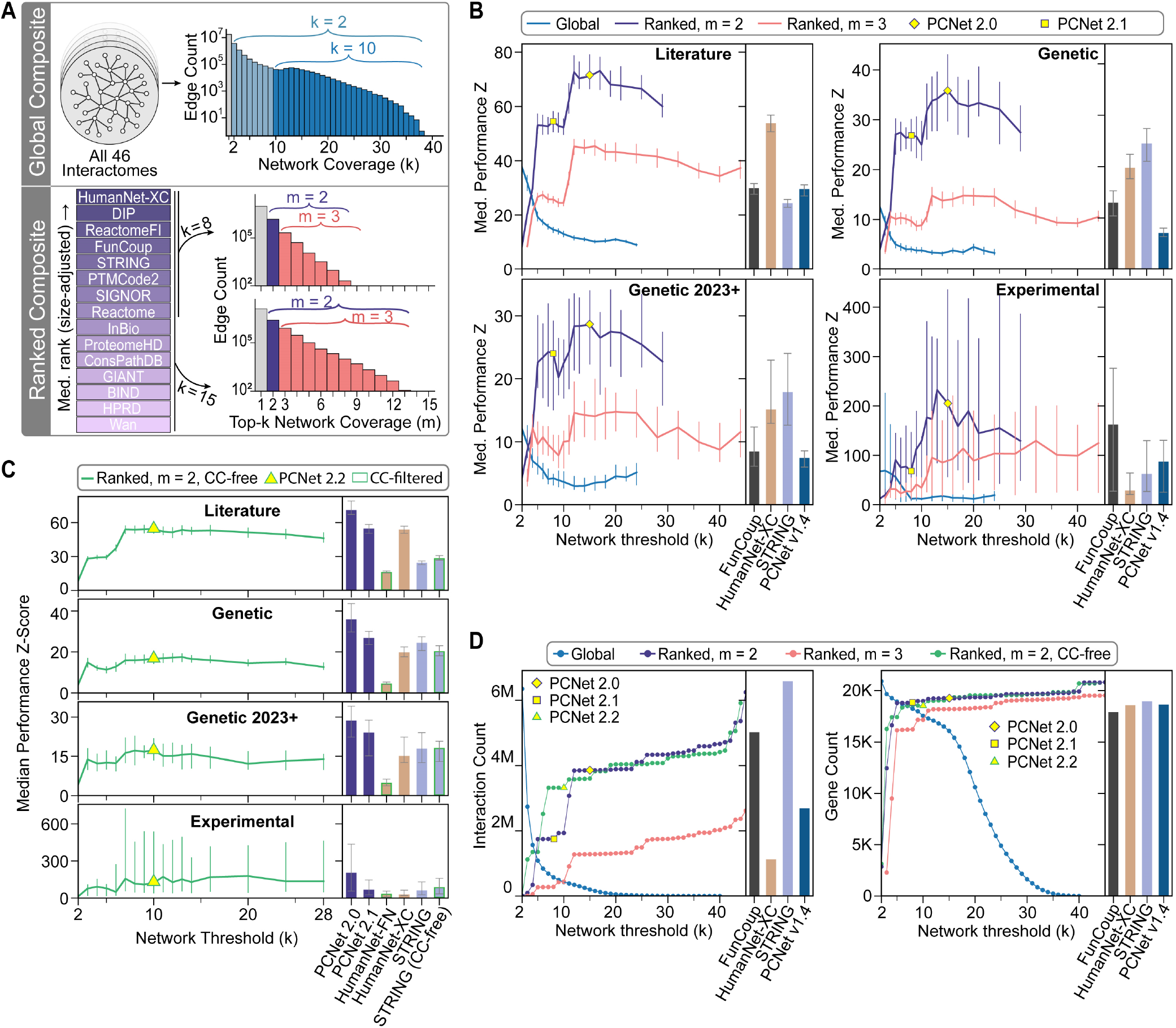
Definition and evaluation of Parsimonious Composite Networks (PCNets). A) Schematic representation of approaches and thresholds used to define global and ranked composite networks. Global composites include all interactions present in at least *k* of 46 interactomes. Ranked composites include all interactions present in at least *m* of the top-*k* interactomes. B-C) Gene set recovery performance for global and ranked composite networks across a range of network thresholds with (B) and without (C) co-citation evidence. Error bars show 95% confidence intervals on the median. Yellow points designate PCNets. Bar charts show equivalent results for comparison interactomes. In (B), results are compared to top-performing individual networks and PCNet v1.4. In (C), results are compared to full and citation-free versions of HumanNet and STRING, as well as PCNet 2.0 and PCNet 2.1 from the present analysis. D) Number of interactions and genes in global and ranked composite networks across a range of network thresholds, compared to top-performing individual interactomes and PCNet v1.4.

Gene set recovery evaluation showed that the performance of global composite networks steadily decreased as the interactome coverage threshold became stricter (Figure 4B). For ranked composites, performance increased as more interactomes were considered, up until 10-15 networks, after which it began to decay. The ranked composite method requiring interaction support from two top-ranked networks showed the best overall gene set recovery performance of all approaches. We also evaluated “co-citation-free” ranked composite networks to mitigate the possible confounding effects of co-citation (CC) interactions. While less powerful than their CC-inclusive counterparts, these CC-free consensus networks still generated strong gene set recovery performance results (Figure 4C).

From these results, we defined an updated set of Parsimonious Composite Networks (PCNets), balancing performance and parsimony. First, we defined the best-performing ranked composite (top 15 interactomes, 3.85M interactions) as PCNet 2.0. While larger than PCNet 1.0, PCNet 2.0 is smaller than many component interactomes, such as STRING^41^ and FunCoup^46^ (Figure 4D). For situations with computational constraints, we also defined the smaller PCNet 2.1 (top 8 interactomes, 1.75M interactions), and for a CC-free alternative, we defined PCNet 2.2 (top 10 CC-free interactomes, 3.32M interactions). These consensus networks are publicly available via NDEx^22^ (ndexbio.org) and include details of supporting interactomes.

### Evaluation and in silico validation of predicted interactions

Finally, we leveraged advances in edge prediction algorithms^19^ and AlphaFold-based modeling^20^ to evaluate the interaction prediction performance of the panel of interactomes. Using 10-fold cross-validation with L3^12^ (paths of length 3) and MPS(T)^19,47^ (Maximum similarity, Preferential attachment Score) algorithms, we predicted interactions for each interactome, excluding interactomes with greater than 1.5M interactions due to computational constraints. The predicted interactions were assessed against held-out and gold-standard interactions. We defined gold-standard physical interactions from multimeric protein complexes recorded in the Comprehensive Resource of Mammalian Protein Complexes (CORUM) database^48^ and pathway interactions from the PANTHER knowledgebase^49^ to capture a range of interactome types.

To evaluate interactome performance, we focused on the precision of the most confident predictions. Across all interactomes, prediction precision was higher for held-out interactions than gold standards, and MPS(T) generally produced stronger predictions than L3 (Figure 5A-C). DIP^27^ and Reactome^50^ showed the highest precision for the held-out task, predicting interactions with average precisions at k (P@k) via MPS(T) of 0.57 and 0.55, respectively (Figure 5A). For gold-standard interactions, networks such as Havugimana^51^ (P@k = 0.19, L3) and PTMCode2^52^ (P@k = 0.166, MPS(T)) led the prediction of CORUM interactions (Figure 5B). In contrast, curated interactomes such as SIGNOR^36^ (P@k = 0.05, L3) and Reactome (P@k = 0.04, L3) showed the best performance for predicting PANTHER pathway interactions (Figure 5C). We further observed that smaller interactomes tended to predict interactions with broad support from other interactomes (Figure 5D).

**Figure 5.**
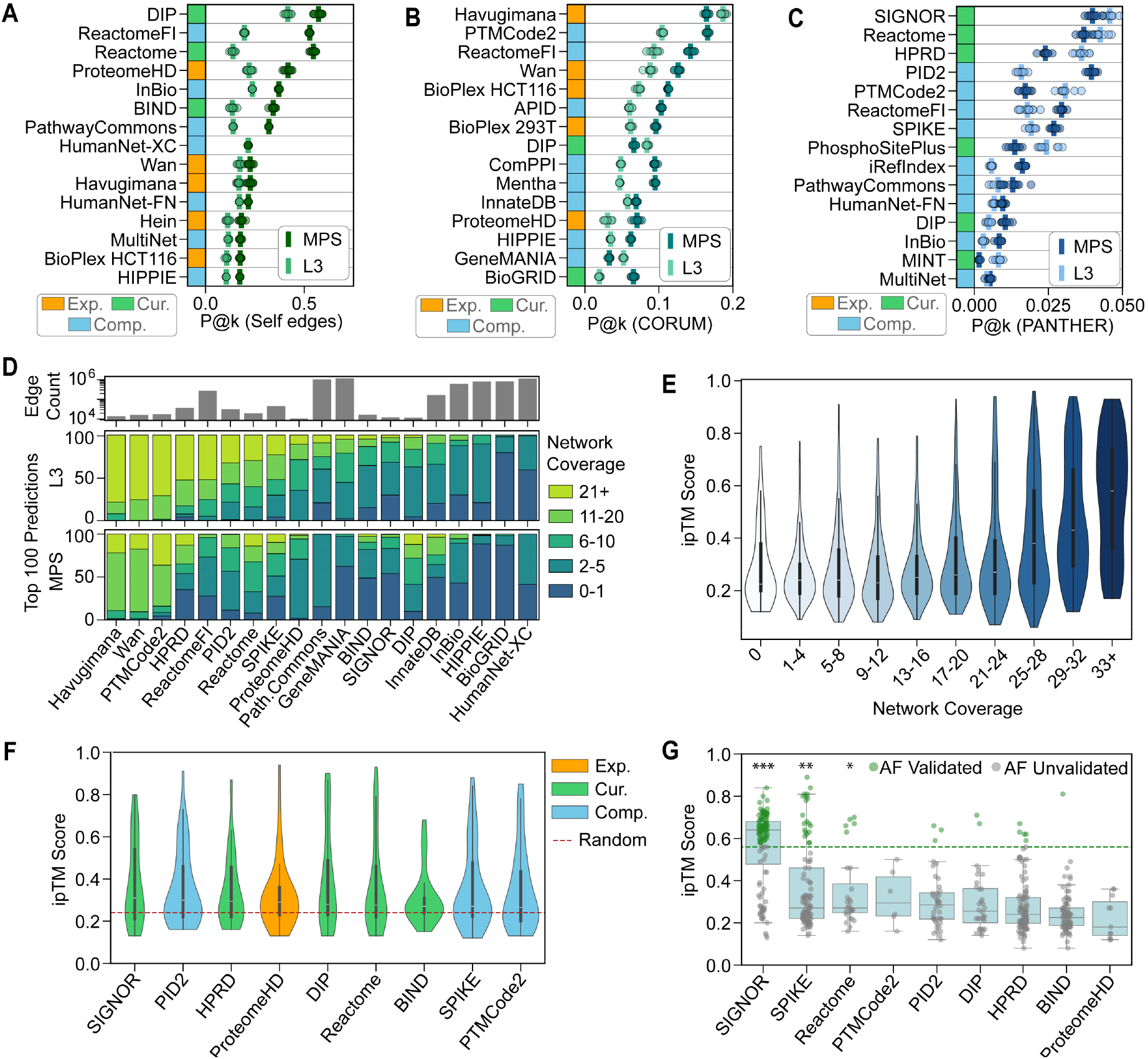
Evaluation of interaction prediction performance with in silico validation. A-C) Interaction prediction evaluation by L3 and MPS(T) using 10-fold cross-validation with test sets defined by (A) held-out self-interactions, (B) CORUM complex interactions, and (C) PANTHER pathway interactions. Mean (bar) and individual (points) precision at k (P@k, k = size of test set) for the top 15 interactomes from each analysis. D) Mean orthogonal network coverage for the top 100 non-self interaction predictions. E) Relationship between AlphaFold-Multimer interface predicted TM-score (ipTM) and interaction network coverage. F) Interactomes enriched for interactions with high interface scores (q < 0.15, Mann-Whitney U-test, BH-corrected), using random samples of 50 network interactions compared to a background distribution of 1779 random protein pairs. G) AlphaFold validation of previously unreported interactions predicted by high-performing interactomes using MPS(T). The significance of predicted interface scores was assessed by the Mann-Whitney U-test against the random background distribution (* q < 0.1, ** q <0.001, *** q<10_–5_, BH-corrected). Validated interactions were defined as protein pairs with an interface score in the 95th percentile (ipTM > 0.56, dashed line) of the random background distribution. See also Supplemental Tables 4-5.

Though not a full substitute for experimental validation, AlphaFold-Multimer^20^ provides an in silico approach to validating potential physical protein interactions. This model builds on the success of AlphaFold^53^, a deep-learning model for predicting the structure of individual proteins. AlphaFold-Multimer specifically trains an AlphaFold model using heterodimer and complex structures from PDB^54^ to enable predictions of multimeric protein interfaces. Applying AlphaFold-Multimer to random selections of interactions with varying interactome coverage, we observed higher interface predicted template modeling (ipTM) scores for interactions present in more than 25 interactomes (Figure 5E), especially for the most supported interactions (> 33 supporting interactomes). We further identified nine interactomes with significantly higher ipTM scores than random protein pairs (Figure 5F). Given these results, we applied AlphaFold-Multimer to validate previously unreported interactions predicted by the nine interactomes. Defining an ipTM score in the 95th percentile (ipTM > 0.56) as a validated interaction, we observed an overall validation rate of 21.5%. The ipTM scores of predicted interactions were significantly higher than random protein pairs for SIGNOR, SPIKE^55^, and Reactome (Figure 5G, Figure S4, FDR < 0.1, Mann Whitney U-test). Interactions predicted by SIGNOR showed particularly high ipTM scores, with 70% being validated, indicating that many of these predictions are likely to represent true physical interactions.

## DISCUSSION

The continued evaluation of interactomes is essential as the range and scope of molecular networks and network databases increase. Here, we have presented the most expansive snapshot of network resources to date (Figure 1), assessing the current state of a diverse range of 46 interactomes, including their features, contents, and performance for discovering disease genes and prioritizing novel interactions. Many of these interactomes now cover nearly all protein-coding genes, though not all biological domains are equally represented (Figure 2F). While some underrepresentation may reflect genes with truly fewer interactions, these results suggest a need for increased focus on less-studied proteins and functions such as transporter and receptor activity.

As interactomes have become more numerous, understanding the utility of different data sources and network structures for network biology applications, such as disease gene prioritization, has become critical^15,40,56^. We found that a small set of networks, including HumanNet, STRING, and FunCoup, consistently produced the strongest disease gene set recovery performance across literature and genetic gene sets (Figure 3B). However, differences in performance between gene set sources highlighted the importance of careful network selection. Performance Z-scores were consistently higher for literature gene sets than genetic gene sets, possibly due to a combination of gene set size and the shared reliance on publications between the literature genesets and the sources used to construct many of the interactomes. However, the interactome rankings remained stable when considering recent genetic findings and experimentally derived gene sets (Figure 3C).

Consistent with previous observations^15,56^, gene set recovery performance was highly correlated with the number of interactions in each network. After adjusting for this size effect, we found that HumanNet-XC and DIP had the highest performance per interaction, indicating that the information in these networks is of very high quality (Figure 3B). In contrast, some high-performing interactomes’ rankings suffered after size adjustment, indicating that they may contain many lower-quality interactions that are nevertheless overcome by the network propagation procedure. We used the size-adjusted interactome rankings to construct high-quality parsimonious composite networks (PCNets), demonstrating even higher gene set recovery performance (Figure 4). These networks, PCNet 2.0, PCNet 2.1, and PCNet 2.2, are publicly available via NDEx.

Further, we developed an interaction prediction evaluation metric to assess the quality of available interactomes. In contrast to gene set recovery results, interaction prediction precision was not driven by interactome size, and interactome rankings changed between different evaluation sets (Figure 5A-C). Therefore, network choice should be carefully considered for predicting potentially novel interactions. Our in silico validation results with AlphaFold-Multimer further reinforce the importance of network selection, with a small subset of interactomes (SIGNOR, SPIKE, and Reactome) predicting interactions with a high AlphaFold-Multimer validation rate (Figure 5G). Our results suggest these networks could be more frequently utilized in interaction prediction studies. It should be noted that the absence of in silico validation for other predicted interactions should not be considered evidence against their existence. While AlphaFold-Multimer provides evidence for possible direct physical interactions, it does not capture the full range of potential gene-gene relationships. Additional in silico and in vitro validation of predicted interactions could build on the interaction prediction evaluation presented here.

In addition to our comprehensive survey, we acknowledge the contributions of other studies to understanding the current state of interactomes. Here, we have used a biological lens to understand the information contained within molecular networks and their performance across a broad range of human disease contexts. This work, therefore, complements detailed studies of network topology^56,57^, re-wiring^10^, and disease and tissue context^58–60^. We anticipate that molecular network data generation and analysis will continue apace, making regular evaluations of available networks essential for optimizing biological discovery. This work provides both an up-to-date benchmarking of available interactomes and a set of refined and expanded tools for ongoing evaluation of biological networks in the future (https://github.com/sarah-n-wright/Network_Evaluation_Tools). While no framework can evaluate all possible interactomes for all applications, we hope this work will serve as a broad guide for network selection, and we welcome the continued development of complementary approaches.

## ACKNOWLEDGEMENTS

This work was supported by the National Institutes of Health under awards U24 HG012107, U24 CA269436, P41 GM103504, and P50 DA037844. The Genotype-Tissue Expression (GTEx) Project was supported by the <u>Common Fund</u> of the Office of the Director of the National Institutes of Health, and by NCI, NHGRI, NHLBI, NIDA, NIMH, and NINDS. The data used for the analyses described in this manuscript were obtained from: the GTEx Portal on 07/11/2023.

We would also like to acknowledge all of the labs and consortia involved in the development and maintenance of all the interactomes we used in this study: Agile Protein Interactomes DataServer (APID), Biomolecular Interaction Network Database (BIND), Biological General Repository for Interaction Datasets (BioGRID), Biophysical Interactions of ORFeome-based comPLEXes network (BioPlex), Compartmentalized Protein-Protein Interaction Database (ComPPI), ConsensusPathDB, Database of Interacting Proteins (DIP), FunCoup, GeneMANIA, Genome-scale Integrated Analysis of gene Networks in Tissues (GIANT, now HumanBase), High-quality INTeractomes (HINT), Human Integrated Protein-Protein Interaction rEference (HIPPIE), Human Protein Reference Database (HPRD), HumanNet, Human Protein Complex MAP (hu.MAP), The Human Reference Interactome (HuRI, formerly the Human Interactome Project), Integrated Interactions Database (IID), ZS (InBioMap, formerly Intomics), iRefIndex, InnateDB, IntAct, MatrixDB, Mentha, Molecular INTeraction database (MINT), MultiNet, Pathway Commons, Pathway Interaction Database (PID), Human Protein-Protein Interaction Prediction (PIPs), PhosphoSitePlus®, Predicted Protein-Protein Interactions (PrePPI), PROPER, ProteomeHD, PTMCode2, Reactome (and Reactome Functional Interactions), SignaLink, The SIGnaling Network Open Resource (SIGNOR), SPIKE, Search Tool for Recurring Instances of Neighboring Genes (STRING), TFLink, the International Molecular Exchange (IMEx) consortium, and the Network Data Exchange (NDEx).

## AUTHOR CONTRIBUTIONS

Conceptualization, SNW, DP, and TI; Methodology, SNW, DP, and TI; Software, SNW, CC, and LVS; Analysis, SNW, SC, and LVS; Manuscript Preparation, SNW, RTP, DP, and TI; Data Curation, SNW, SC, and RTP; Funding Acquisition, DP and TI.

## AVAILABILITY OF DATA AND MATERIALS

All code used for analysis and data visualization is freely available in public repositories. All original code is publicly available at GitHub: https://github.com/sarah-n-wright/Network_Evaluation_Tools and will be deposited at Zenodo upon publication. Links to all source data are provided in Supplemental Table 1.

All networks described in this manuscript have been made publicly available via the Network Data Exchange (NDEx). NDEx is an open-source, publicly available software infrastructure that facilitates the storage, exchange, visualization, and publication of network models and data among scientists. Thanks to its full integration with the Cytoscape desktop application, users can employ NDEx to import, export, and analyze biological networks using the large variety of tools and applications available in the Cytoscape ecosystem. The platform’s key advantages include fostering collaborative research through shared networks, enabling the reproducibility of scientific findings, and promoting the discovery of new biological insights by integrating disparate data types into comprehensive network models. NDEx thus offers a centralized resource to enhance the utility and accessibility of network-based data and analyses in elucidating complex biological systems. To learn more about NDEx, please review the FAQ page at www.ndexbio.org.

The complete suite of networks is available in the dedicated PCNet 2.0 NDEx account at https://www.ndexbio.org/index.html#/user/bae4da70-e22d-11ee-9621-005056ae23aa.

Digital Object Identifiers (DOI) allow direct access to the set containing all source networks (https://doi.org/10.18119/N95C9J), as well as the 3 Parsimonious Composite Networks: PCNet2.0 (https://doi.org/10.18119/N9JP5J), PCNet2.1 (https://doi.org/10.18119/N9DW40), and PCNet2.2 (https://doi.org/10.18119/N9960N).

## METHODS

### DATA COLLECTION AND PROCESSING

#### Collection and standardization of interaction data

Where possible, all networks were downloaded from primary sources (Supplemental Table 1). DIP^61^ and BIND^62^ were downloaded from PathwayCommons v12 ^63^, and PCNet 1.4^15^ was downloaded from NDEx^21,22^ (ndexbio.org). All non-human interactions were excluded, except in cases where the authors used orthologous interactions to enhance human networks (Figure 1A). Duplicated and self-interactions were removed, and non-binary interactions were binarized by defining edges between all pairs of genes. All interactions were treated as undirected. The GeneMANIA^45^, GIANT^58^, and hu.MAP 2.0^64^ interactomes contained > 15M interactions. Therefore, we filtered these networks to the top 10% of interactions using the provided interaction scores. The STRING co-citation-free (CC-free) network was defined by excluding interactions supported solely by the ‘textmining’ channel.

#### Collation of network metadata

Network dependencies and interaction types were collated from publicly available information based on definitions in Supplemental Table 6. A network dependency between a target and a source network was defined as a relationship where interactions from the source network were directly incorporated into the target network. The use of a network to train, prioritize, or score interactions was not considered a dependency. Where available, PMIDs associated with interactions were used to identify experimental sources.

#### Standardization of gene identifiers

We mapped all gene identifiers to NCBI Gene IDs by utilizing APIs from MyGeneInfo^65,66^, UniProt^67^, HGNC^33^, and Ensembl^68^. Input identifiers were updated using the most relevant database to address out-of-date identifiers, and then converted to NCBI Gene IDs. For HGNC Gene Symbols, previously approved symbols were prioritized over alias symbols. The performance of our gene mapping pipeline was compared to the use of MyGeneInfo alone. For dissemination of networks we retained Gene IDs as the primary identifiers while reporting the current approved HGNC gene symbols (as of April 1, 2024) for ease of use.

#### Collation and processing of gene metadata

We sourced gene and protein annotation features from HGNC^33^ (chromosome, locus type, locus group) and NCBI^69^ (citation count). The set of protein-coding genes was defined by the HGNC Locus Group ‘protein-coding gene.’ Functional Gene Ontology (GO) associations were downloaded from NCBI using goatools^70^ on March 29, 2023.

To assess mRNA expression patterns, we sourced median gene-level TPM by tissue from GTEx (v8, August 2019). For genes with multiple transcripts, we consolidated the values within each tissue. We took the mean of all transcripts if all values had a similar magnitude (all TPM observations above or below 10^-4^); otherwise, we took the maximum.

For protein expression, we utilized normal tissue expression values by tissue reported by the Human Protein Atlas^34,35^ (HPA) v23 (https://www.proteinatlas.org/download/normal_tissue.tsv.zip). We excluded entries with ‘Uncertain’ reliability and tissues with fewer than 1000 observations. We consolidated values for each gene by taking the mean of all associated protein expression values. Categorical expression levels were converted to numerical values (0=Not detected, 1=Low, 2=Medium, 3=High).

#### Definition of gene and interaction sets

We generated collections of genesets from three sources for the evaluation of gene set recovery performance and converted all gene identifiers to NCBI Gene IDs. First, we obtained literature-curated disease gene sets from DisGeNET^71^, utilizing BEFREE gene-disease associations sourced from the text-mining of MEDLINE abstracts. We retained sets with a maximum of 500 genes and at least 5 genes in every interactome. Second, we created genetic gene sets from genome-wide association studies (GWAS) via the GWAS Catalog^39^. From the full download (https://www.ebi.ac.uk/gwas/api/search/downloads/alternative), we extracted all SNPs mapped to an NCBI Gene ID with a significant association (p-value < 5e-8) to a phenotype in the Experimental Factor Ontology (EFO). We created gene sets for all phenotypes with fewer than 500 gene associations across all studies. In addition, we create a collection of recent gene sets using only GWAS with a publication date after July 27, 2023, thus postdating the latest update of interactomes incorporating co-citation interactions. Lastly, we identified gene sets generated from experimental data and published after January 1, 2024. Sets in this experimental collection were defined primarily from differential expression studies. To analyze each interactome, we selected genesets with a minimum of 20 (literature, genetic) or 10 (recent genetic, experimental) genes and a maximum of 500 genes. Gold-standard complex and pathway interactions were sourced from CORUM^48^, KEGG^72^, and PANTHER^49^ via PathwayCommons^63^. Interactions were standardized using the same procedure as for interactomes above to generate interaction lists. All gene sets are available in Supplemental Table 7.

#### Creation of interaction-type-specific networks

We defined interaction-type-specific from HumanNet v3^40^ and GeneMANIA^45^. From HumanNet, we utilized HS-CC (co-citation), HS-CX (co-expression), HS-DB (pathway), HS-DP (domain), HS-GI (genetic interaction), HS-PG (phylogenetic similarity), and HS-PI (physical). All networks were downloaded directly and standardized per our pipeline outlined above. From GeneMANIA, we downloaded source files from https://genemania.org/data/current/Homo_sapiens/ based on assigned types (‘Co-expression,’ ‘Co-localization,’ ‘Genetic_Interactions,’ ‘Pathway,’ ‘Physical_Interactions,’ ‘Predicted,’ ‘Shared_protein_domains’). All interactions within a source were concatenated to form interaction-type-specific networks. GeneMANIA sources for genetic and co-expression interactions report all possible gene pairs with interaction scores. Therefore, we filtered these interaction-type-specific networks to the top 10% of genetic interactions and the top 3% of co-expression interactions based on interaction scores. We assigned HS-PG and GeneMANIA-Predicted to the category ‘Orthology’ as both utilized information from non-human species.

#### Creation of composite/consensus networks

First, we defined ‘Global Composite” networks *G*_k_ based on the frequency of each interaction across all interactomes, excluding HumanNet-FN (which is fully contained in HumanNet-XC). For global composite networks, we defined the network threshold *k* as the minimum number of supporting networks for each interaction. For example, *G*_k=2_ includes all interactions occurring in at least two interactomes. Formally, let *i* be an undirected interaction, *n* be the number of interactomes, *I*_i,1_, *I*_i,2_,…, *I*_i,n_ indicate whether an interaction *i* is present in each interactome, and *k* be the network threshold. Then:

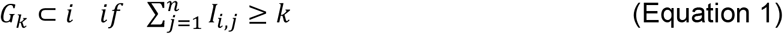

Secondly, we define ‘Ranked Composite’ networks 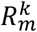 based on the frequency of each interaction in the top-*k* best-performing networks. The best-performing networks were defined based on the average rank of size-adjusted performance across all literature and genetic gene sets. For ranked composite networks, we define the network threshold *k* as the number of top-performing networks considered. We define a second threshold *m* to represent the threshold on the number of supporting networks for each interaction. For example, 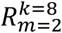 includes all interactions present in at least two of the top eight interactomes. Formally, let *i* be an undirected interaction, *n* be the total number of interactomes sorted by rank, *I*_i,1_, *I*_i,2_,…, *I*_i,n_ indicate whether an interaction *i* is present in each interactome, *k* be the network threshold (number of top networks considered), and *m* be the network support threshold. Then:

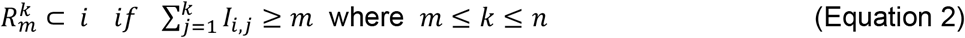

#### Deposition of interactomes to NDEx

All standardized source networks were uploaded to the NDEx network set “State of the Interactomes: source networks” via the NDEx2 Python Client^22^ v3.5.0. All genes were indexed by NCBI Gene IDs and annotated with current approved HGNC Symbols as of April 2, 2024. The original gene identifiers used by the source database were maintained, as were additional edge annotations where available, such as PubMed IDs, interaction type, or detection method. PCNet 2.0, PCNet 2.1, and PCNet 2.2 were similarly uploaded via the NDEx2 Python Client. Genes were annotated with the current approved HGNC Symbols, and interactions were annotated with the number of supporting interactomes and a list of those supporting interactomes.

#### REPRESENTATION ANALYSIS

Spearman correlations between gene features and network coverage were calculated using scipy^73^. For mRNA and protein expression, genes were binned into 20 expression percentiles based on the mean expression across all tissues. Due to the number of genes with near-zero average mRNA expression, the lowest 20th percentile was treated as a single bin for mRNA expression. Citation counts were adjusted for mean mRNA expression levels using a log-log ordinary least squares linear regression via statsmodels^74^.

To calculate the representation of biological processes, molecular functions, and cellular compartments, we calculated gene set enrichment via Fisher’s Exact Test for GO Slim terms (https://current.geneontology.org/ontology/subsets/goslim_generic.obo). From the GO Slim ontology, we selected terms with 100-4000 associated genes and excluded any terms that were parents of other terms in the set, resulting in a set of 93 terms. The background for enrichment analysis was set to all protein-coding genes (defined by HGNC^32^) with at least one GO association, and all genes associated with a term were also considered to be associated with all parents of that term.

To assess the relationships between gene annotation features and the presence of interactions across all networks, we assessed the number of interactions between genes with varying citation count, mRNA expression, protein expression, and chromosome number. Each interaction GeneA-GeneB present in at least one interactome was assigned two bins: BinA corresponding to the annotation value for GeneA and BinB corresponding to the annotation for GeneB. Across all unique interactions from all 46 interactomes, the number of interactions between all combinations of BinA and BinB (*n*_AB_) was calculated and normalized based on the number of possible interactions between genes in BinA (*A*) and BinB (*B*) to give the interaction fraction *f*_AB_:

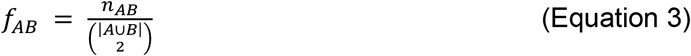

The same process was applied for GO gene-term associations for the set of 93 GO Slim terms considered, allowing for multiple associations per gene.

### GENE SET RECOVERY EVALUATION

#### Network Propagation with Subsampling

Gene set recovery via network propagation was implemented following the procedure previously established^15^ for each interactome and gene set. First, we took the intersection of interactome and gene set genes. Then, we randomly sub-sampled a proportion *p* without replacement from this intersection as our seed genes, with the remaining portion of the gene set becoming the held-out set. This group of seed genes was propagated through the network using a random walk with restart model^75^, utilizing the closed-form solution^76^:

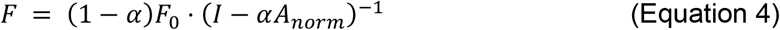

Where *F* is the vector of propagation scores for all genes, *α* is the network propagation constant, *F*_O_ is a binary vector representing each gene’s initial seed gene status, and *A*_norm_ is the degree-normalized adjacency matric of the interactome.

Defining the held-out set *h* as the true positives, we ranked all interactome genes by propagation score and calculated the area under the precision-recall curve (AUPRC). The sampling, propagation, and AUPRC calculations were repeated for 50 samples and then averaged to give a mean AUPRC for each gene set - interactome pair. To assess the significance of the observed AUPRC values, we constructed 50 randomized versions of each interactome via degree-matched edge shuffling and repeated the analysis for each gene set to give a null distribution of mean AUPRC values.

We defined the ‘Performance’ score as the robust Z-statistic^77^ between the observed and null mean AUPRC values. We defined the ‘Performance Gain’ as the difference between the observed mean AUPRC and the median AUPRC of the associated null networks, normalized by the median AUPRC of the associated null networks. Additionally, we calculated a size-adjusted performance metric by fitting a linear regression model for each gene set between the log_10_ interaction count and the performance score. The residual for each interactome was taken as the size-adjusted performance. All gene set recovery results are available in Supplemental Table 3.

#### HParameter Optimization for Gene Set Recovery

The two parameters *α* (network propagation constant) and *p* (subsampling proportion) were optimized for each interactome and gene set using the 50 MSigDB Hallmark pathways^78^ as an independent source of gene sets. We compared gene set recovery performance across intervals of *α* ∈ {0.2, 0.3, …,0.9} and *p* ∈ {0.1, 0.2, … 0.8} for all interactomes and MigDB gene sets (Figure S5A, Supplemental Table 8). We performed a linear regression of mean performance using Huber Regression^79^ via sklearn^80^ with *α, p*, network size (normalized log10 interaction count) and gene set coverage (normalized size of intersection between gene set and interactome genes) as features. This analysis identified that the subsampling parameter *p* had a stronger influence on performance than the propagation constant *α*. To model the optimal parameters, we first calculated the average optimal parameter values for each interactome-gene set pair by taking the mean of the top five performing parameter sets. We then fit the average optimal subsampling parameter to the normalized network size and gene set coverage (Figure S5B). The resulting formula was used to set the subsampling parameter for each interactome-gene set pair:

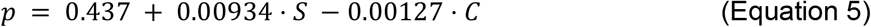

Where *S* is the log10 network size (interaction count), and *C* is the gene set coverage. We restrict the value to 0.1 < *p* < 0.8. Next, we fit the average optimal propagation constant for each interactome to the normalized network size, and the average subsampling parameter and gene set coverage across all MSigDB gene sets (Figure S5C). The resulting formula was used to set the propagation constant for each interactome:

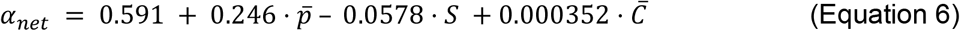

Where 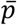 is the mean subsampling parameter across all gene sets for the interactome, *S* is the log10 network size (interaction count), and 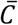 is the mean gene set coverage across all gene sets for the interactome. We restrict the value to 0.2 < *α*_*net*_ < 0.9. For type-specific, global composite, and ranked composite networks, we set the subsampling parameter using Equation 5, and we set the propagation constant by taking the mean optimal value (*α*_*net*_ = 0.64) across all interactomes.

#### Network Performance Rankings

To compare the performance of interactomes, we first calculated a centralized rank from the performance Z-scores for each gene set across all interactomes and then took the mean centralized rank across all gene sets for each interactome. Our centralized rank formulation accounts for not all interactomes having sufficient gene set coverage of every gene set (Figure S3A). For each gene set, we ranked the performance scores of all *m* successfully evaluated interactomes and centralized this rank to give values evenly distributed around zero. Thus, for a given gene set:

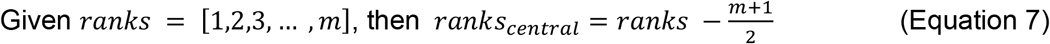

As is typical for rank measures, the lower the centralized rank, the better the relative performance of an interactome. In this way, a negative centralized rank reflects an interactome in the top 50% of interactomes assessed, and a positive centralized rank reflects an interactome in the bottom 50% of interactomes assessed. This measure ensures the most weight is given to results for gene sets assessed with all interactomes while still utilizing results from smaller gene sets that could only be assessed with a smaller number of large networks.

Centralized ranks were calculated separately for literature, genetic, and experimental gene set sources for both raw and size-adjusted performance scores. The mean centralized rank of size-adjusted performance across literature and genetic gene sets was used to rank all interactomes for consideration in creating ranked composite networks. PCNet 2.0, PCNet 2.1, and PCNet 2.2 were selected based on the mean centralized rank across literature and genetic genesets from all global and ranked composite networks (or all co-citation-free global and ranked composite networks). For PCNet 2.1, we additionally restricted the maximum number of interactions to two million.

### INTERACTION PREDICTION AND EVALUATION

#### Interaction prediction performance

We implemented two edge prediction algorithms to assess interaction prediction accuracy: L3^12^ and MPS(T)^19,47^. The L3 method ranks predicted edges based on the number of paths of length three between two nodes, normalized by the degree of the intermediate nodes. The MPS(T) method utilizes two measures of topological similarity (Maximum similarity and Preferential attachment Score) based on the hypothesis that the probability of two proteins interacting is proportional to the similarity between one protein and the most similar interactors of the other protein. Implementations of the L3 and MPS(T) algorithms were sourced from Github: L3 v.1.0.2 (https://github.com/kpisti/L3) and MPS(T) (https://github.com/spxuw/PPI-Prediction-Project/). We standardized the prediction evaluation for both methods.

We first measured the performance of interaction prediction via a 10-fold cross-validation procedure by performing prediction with 90% of network edges and assessing the recovery of the 10% of held-out edges. We used a precision at k (P@k) metric to calculate interaction prediction performance. This metric focused the evaluation on the top predicted interactions, which are typically of most interest to researchers, and allowed rapid calculation by avoiding consideration of up to 400M interactions for the largest networks. To create a fair benchmark between interactomes, we set k to be the size of the test set (e.g., 10% of interactome size for held-out edges).

In addition to the prediction of held-out edges, we assessed the ability of networks to predict gold-standard interaction sets defined by CORUM^48^ and PANTHER^49^. Each of the 10 folds created for cross-validation was used for prediction, with performance calculated again as P@k for the gold-standard edge sets, excluding any gold-standard edges already present in the network. In some cases, network structure and edge removal led to genes with degree zero. Because the prediction algorithms utilize network topology, predicting interactions to genes of degree zero is impossible. Therefore, any such interactions in the held-out or gold-standard sets were excluded from performance calculations. All interaction prediction results are available in Supplemental Table 4.

#### Computational validation of predicted interactions with AlphaFold-Multimer

Validation of predicted interactions was performed using AlphaFold-Multimer^20^ using ColabFold^81^ v1.5.5 (https://pypi.org/project/colabfold/) via localcolabfold (https://github.com/YoshitakaMo/localcolabfold). We set parameters --num-recycle 3 --model-type alphafold2_multimer_v3, with all other parameters set to defaults. For all analyses, interactions were first subset to those between genes that could be mapped to a protein sequence in UniProt^67^. Protein sequences were sourced from UniProt on February 5, 2024.

From the interface predicted template modeling (*ipTM*) and predicted template modeling (*pTM*) scores generated by AlphaFold-Multimer, we took the results from the model with the highest confidence^20^ defined by:

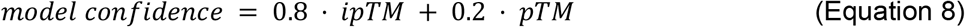

For validation with AlphaFold2-Multimer, we first established a baseline relationship between the network coverage of an interaction and its ipTM score from multimeric structural prediction. To construct this baseline, we randomly selected groups of 50 interactions with network coverage between 1-33 interactomes and 50 interactions with network coverage ≥ 34.

Secondly, we assessed the validation rate for reported interactions within each interactome. We evaluated 50 random protein pairs from each network and compared them to a background distribution of ipTM scores for 1779 random protein pairs selected independently of this study. Statistical testing was performed using a Mann-Whitney U-test.

Lastly, we defined the nine networks with significantly higher ipTM scores than random as candidates for AlphaFold-Multimer Validation. For each of these interactomes, we extracted the union of the top 100 predictions across all folds and defined ‘previously unreported’ protein pairs as those with no reported interactions in any of the 46 interactomes studied. We modeled the interactions between these protein pairs and compared the resulting ipTM scores to the background distribution. We defined a validated interaction as an ipTM score > 0.56, corresponding to the top 5% of the background distribution.

All AlphaFold-Multimer evaluations were conducted using NVIDIA Tesla V100 GPUs operating under Rocky Linux release 8.9. Each evaluation was allocated one CPU and 14 GB of memory and was subject to a 5-hour time limit. Of all protein pairs, 95% of network coverage pairs, 93% of network-specific pairs, and 97% of previously unreported pairs were successfully evaluated. Summarized results of AlphaFold-Multimer modeling are available in Supplemental Table 5.

## SUPPLEMENTAL FIGURES

**Supplemental Figure 1.**
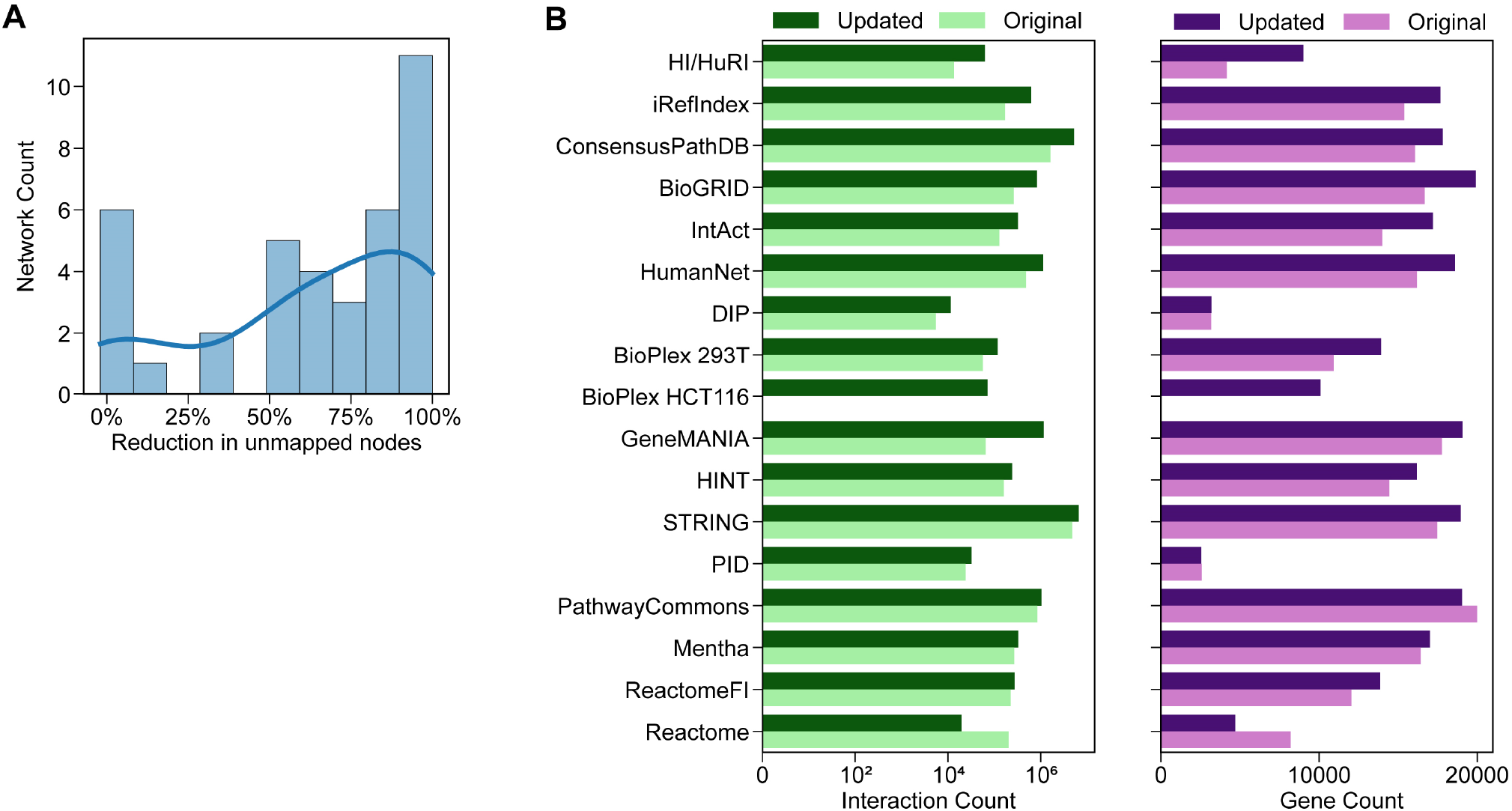
Comparison between original and updated networks and pipeline. A) Change in the number of unmapped interactors using our updated pipeline compared to MyGeneInfo alone. B) Interaction and gene counts for networks updated between Huang et al., 2018 and the present study. Updated networks are those where the corresponding database has been updated or where our source of the network was updated.

**Supplemental Figure 2.**
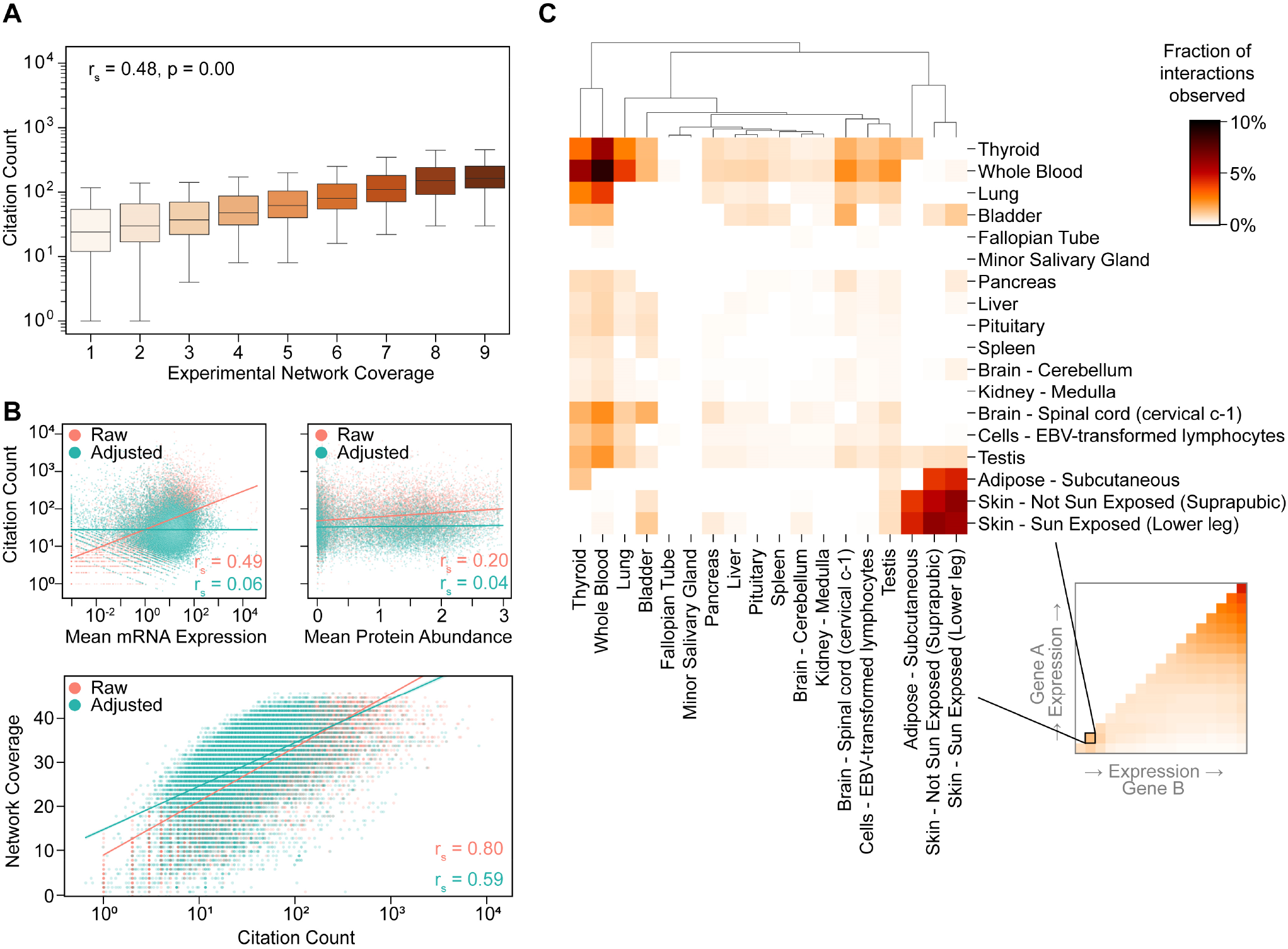
Targeted analysis of gene and interaction annotations. A) Spearman correlation between citation count and network coverage in the nine experimental interactomes. B) Citation counts adjusted for mean gene mRNA expression levels using log-log ordinary least-squares and Spearman correlations to mRNA expression, protein abundance, and network coverage. C) Interaction density of select tissue-specific genes. Genes were taken from the 20-25th percentile of mean mRNA expression (n=1740) and filtered to those with non-zero expression in a maximum of 2 tissues (1400/1740). Tissues with at least one gene having a reported interaction are shown.

**Supplemental Figure 3.**
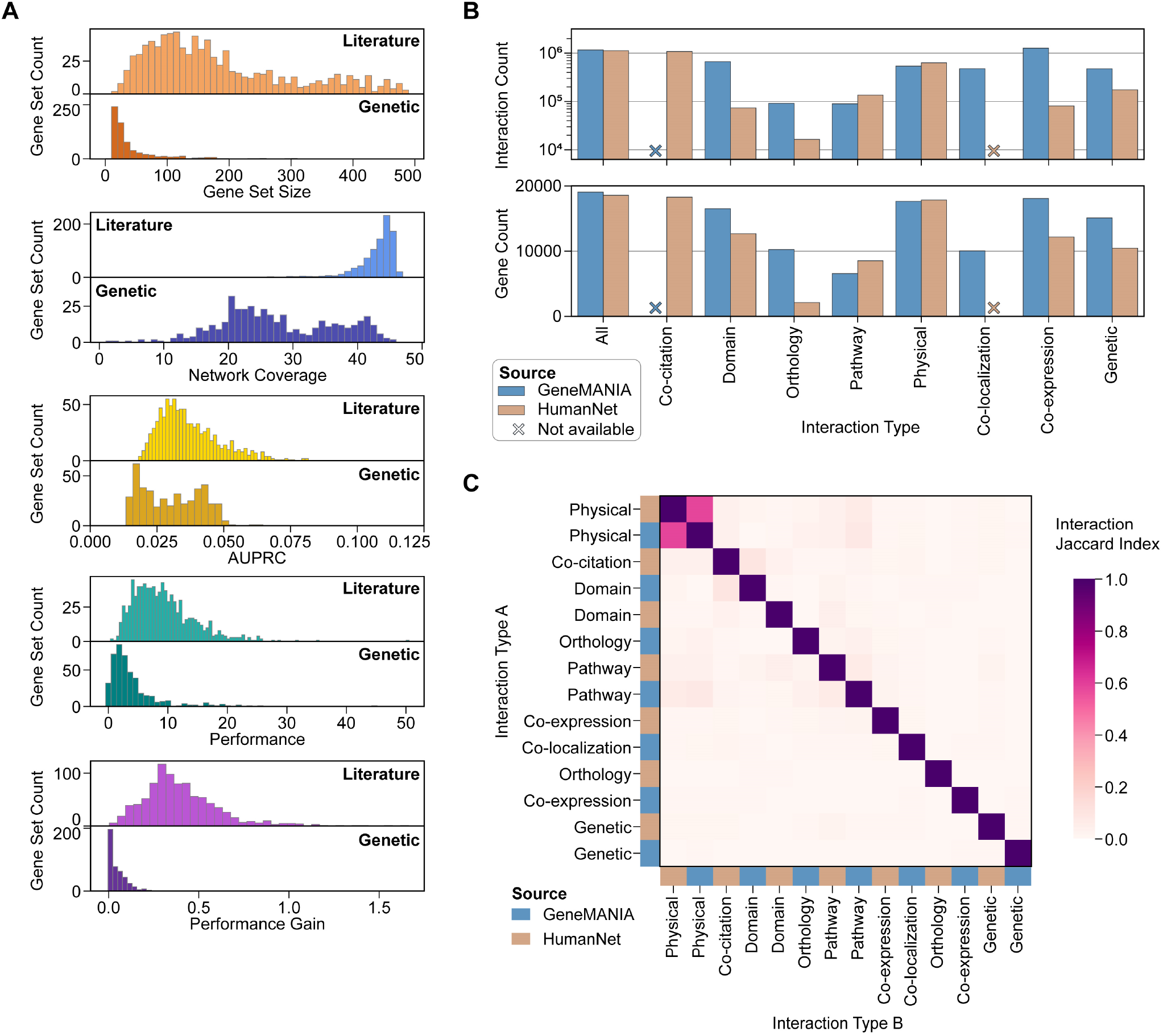
Descriptive statistics of gene set recovery performance and type-specific networks. A) Distributions of gene set statistics and performance metrics. Gene set size: number of unique genes in each set after mapping to NCBI Gene IDs. Network Coverage: number of networks containing at least 20 set genes. AUPRC, Performance, Performance Gain metrics: mean metric of a gene set across all networks containing at least 20 set genes. B) Sizes of type-specific interactomes defined from HumanNet v3 and GeneMANIA. Crosses indicate that a network could not be defined from available data. Interaction type ‘All’ refers to the HumanNet-XC and GeneMANIA (top 90%) networks used in the primary analysis. C) Clustered interaction similarities of type-specific networks, measured by the Jaccard Index of network interactions.

**Supplemental Figure 4.**
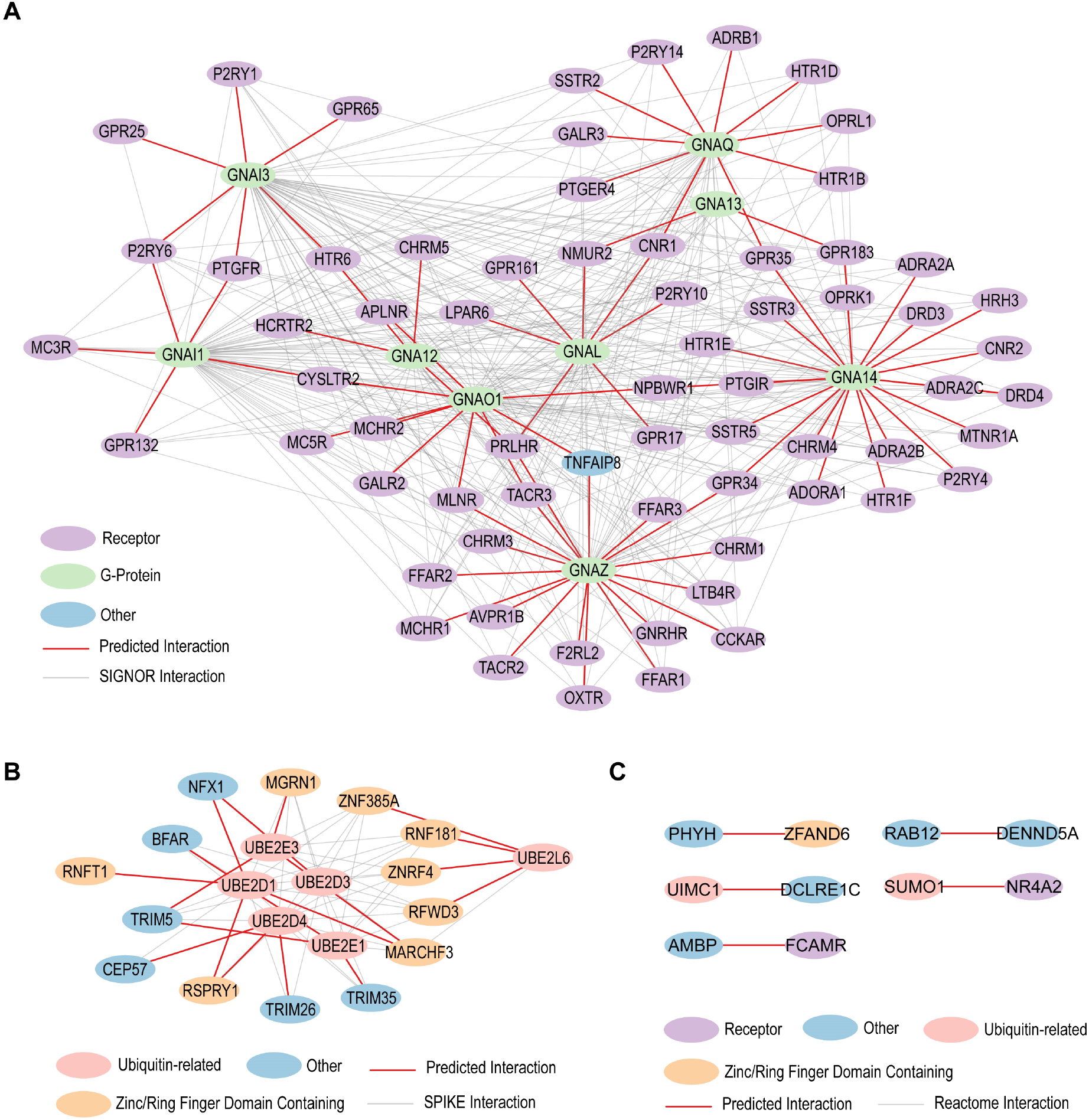
Subnetworks of AlphaFold-Multimer validated predicted interactions. Subnetworks are shown for (A) SIGNOR, (B) SPIKE, and (C) Reactome predicted interactions. Node color indicates broad protein type and edge color indicates prediction status.

**Supplemental Figure 5.**
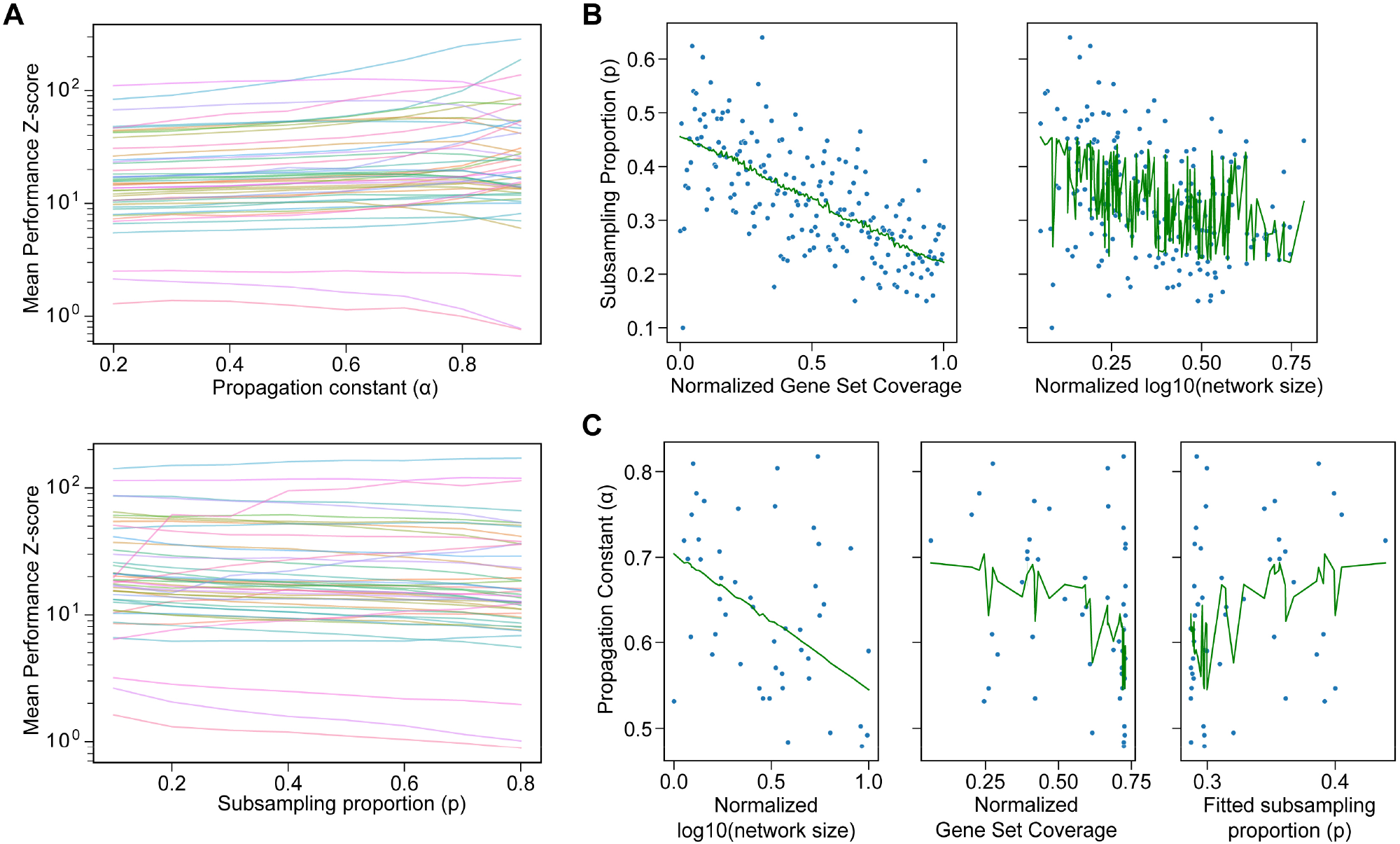
Optimization of gene set recovery parameters. A) Mean performance Z-score for all interactomes across a range of propagation constants (top) and subsampling parameters (bottom). B) Linear fit of gene set coverage and network size to the optimal subsampling parameter. C) Linear fit of network size, average gene set coverage, and average fitted subsampling parameter to the optimal propagation constant. Network size measured as the number of unique interactions within each network.

