## Supplemental Table 1 for "State of the Interactomes: an evaluation of molecular networks for generating biological insights"

**Supplemental Table 1. Interactome source, version, and citation information.** Any networks without version numbers were versioned based on the latest source update or publication date, or assigned version = 1 for interactomes with a single version.

| Name | Version | Source Link | Source Website | Refs |
|---|---|---|---|---|
| APID | 2021 | http://cicblade.dep.usal.es:8080/APID/init.action#tabr1 | http://cicblade.dep.usal.es:8080/APID/init.action | [1,2] |
| BIND | Pathway Commons v12 | https://www.pathwaycommons.org/archives/PC2/v12/PathwayCommons12.bind.hgnc.txt.gz | https://www.pathwaycommons.org/pc2/ | [3] |
| BioGRID | 227 | https://downloads.thebiogrid.org/Download/BioGRID/Release-Archive/BIOGRID-4.4.227/BIOGRID-ORGANISM-4.4.227.tab3.zip | http://thebiogrid.org | [4] |
| BioPlex 293T | 3 | https://bioplex.hms.harvard.edu/data/BioPlex_293T_Network_10K_Dec_2019.tsv | http://bioplex.hms.harvard.edu | [5] |
| BioPlex HCT116 | 3 | https://bioplex.hms.harvard.edu/data/BioPlex_HCT116_Network_5.5K_Dec_2019.tsv | http://bioplex.hms.harvard.edu | [5] |
| ComPPI | 2.1.1 | https://comppi.linkgroup.hu/downloads; options: PPI, H. sapiens | https://comppi.linkgroup.hu/ | [6] |
| ConsensusPathDB | 35 | http://cpdb.molgen.mpg.de/download/ConsensusPathDB_human_PPI.gz | http://cpdb.molgen.mpg.de/ | [7] |
| DIP | Pathway Commons v12 | https://www.pathwaycommons.org/archives/PC2/v12/PathwayCommons12.dip.hgnc.txt.gz | https://www.pathwaycommons.org/pc2/ | [8] |
| FunCoup | 5 | https://funcoup.org/downloads/download.action?type=network&instanceID=24480085&fileName=FC5.0_H.sapiens_full.gz | https://funcoup.org/ | [9] |
| GeneMANIA | 2021 | http://genemania.org/data/current/Homo_sapiens.COMBINED/COMBINED.DEFAULT_NETWORKS.BP_COMBINING.txt | https://genemania.org/data | [10] |
| GIANT | 1 | https://s3-us-west-2.amazonaws.com/humanbase/networks/global_top.gz | https://hb.flatironinstitute.org/ | [11] |
| Havugimana | 1 | Table S2 | http://human.med.utoronto.ca/ | [12] |
| Hein | 1 | https://www.ebi.ac.uk/intact/search?query=IM-24272&interactorSpeciesFilter=Homo%20sapiens,Mus%20musculus,Saccharomyces%20cerevisiae | https://www.ebi.ac.uk/intact/ | [13] |
| HINT | v4 | http://hint.yulab.org/download/ | http://hint.yulab.org/ | [14] |
| HIPPIE | 2.3 | http://cbdm-01.zdv.uni-mainz.de/~mschaefer/hippie/HIPPIE-current.mitab.txt | http://cbdm-01.zdv.uni-mainz.de/~mschaefer/hippie/information.php | [15] |
| HPRD | 9 | http://hprd.org/download (HPRD_Release9_041310.tar.gz) | http://hprd.org/ | [16–18] |
| HumanNet-FN | v3 | https://staging2.inetbio.org/humannetv3/networks/HumanNet-FN.tsv | https://staging2.inetbio.org/humannetv3/ | [19] |
| HumanNet-XC | v3 | https://staging2.inetbio.org/humannetv3/networks/HumanNet-XC.tsv | https://staging2.inetbio.org/humannetv3/ | [19] |
| hu.MAP | 2 | http://humap2.proteincomplexes.org/static/downloads/humap2/humap2_ppis_geneid_20200821.pairsWprob.gz | http://humap2.proteincomplexes.org/ | [20] |
| HuRI | HI-union | http://www.interactome-atlas.org/data/HI-union.tsv | http://www.interactome-atlas.org/ | [21,22] |
| IID | 2021-05 | http://iid.ophid.utoronto.ca/static/download/human_annotated_PPIs.txt.gz | http://iid.ophid.utoronto.ca/ | [23] |
| InBio | 2016 | https://zs-revelen.com/download | https://www.zs-revelen.com/ | [24] |
| InnateDB | 5.4 | https://www.innatedb.com/download/interactions/innatedb_all.mitab.gz | https://www.innatedb.com/ | [25–27] |
| IntAct | 245 | https://ftp.ebi.ac.uk/pub/databases/intact/current/psimitab/intact.zip | https://www.ebi.ac.uk/intact/home | [28] |
| iRefIndex | 20 | https://storage.googleapis.com/irefindex-data/archive/release_20.0/psi_mitab/MITAB2.6/9606.mitab.08-28-2023.txt.zip | https://irefindex.vib.be// | [29] |
| MatrixDB | 2019 | http://matrixdb.univ-lyon1.fr/download/matrixdb_FULL.tab.gz | http://matrixdb.univ-lyon1.fr/ | [30] |
| Mentha | 23.11.6 | https://mentha.uniroma2.it/dumps/organisms/all.zip | https://mentha.uniroma2.it/index.php | [31] |
| MINT | 1 | http://www.ebi.ac.uk/Tools/webservices/psicquic/mint/webservices/current/search/query/* | https://mint.bio.uniroma2.it/ | [32] |
| MultiNet | 1 | http://homes.gersteinlab.org/Khurana-PLoSCompBio-2013/Multinet.interactions.network_presence.txt | http://homes.gersteinlab.org/Khurana-PLoSCompBio-2013/ | [33] |
| Pathway Commons | 12 | https://www.pathwaycommons.org/archives/PC2/v12/PathwayCommons12.All.hgnc.txt.gz | pathwaycommons.org | [34] |

| Name | Version | Source Link | Source Website | Refs |
|---|---|---|---|---|
| PhosphositePlus | Oct18_23 | https://www.phosphosite.org/staticDownloads; Kinase_Substrate_Dataset.gz | https://www.phosphosite.org/ | 35 |
| PID | 2 | https://www.ndexbio.org/index.html#/networkset/7bc65b82-2a2f-11ed-ac45-0ac135e8bacf | https://www.ndexbio.org/index.html#/ | 36 |
| PIPs | 1 | https://www.compbio.dundee.ac.uk/www-pips/dbStats.jsp | https://www.compbio.dundee.ac.uk/www-pips/index.jsp | 37 |
| PrePPI | 2023 | https://honiglab.c2b2.columbia.edu/PrePPI/ref/preppi.human_af.interactome.txt.tar.gz | https://honiglab.c2b2.columbia.edu/PrePPI/ | 38 |
| PROPER | 1 | https://genemo.ucsd.edu/proper/PROPER_v1.csv | https://genemo.ucsd.edu/proper/ | 39 |
| ProteomeHD | 1 | https://www.proteomehd.net/download_file/S3 | https://www.proteomehd.net/index | 40 |
| PTMCode2 | 2 | https://ptmcode.embl.de/data/PTMcode2_associations_between_proteins.txt.gz | https://ptmcode.embl.de/ | 41 |
| Reactome | 86 | https://reactome.org/download/current/interactors/reactome.homo_sapiens.interactions.tab-delimited.txt | https://reactome.org/ | 42 |
| ReactomeFI | 2022 | https://reactome.org/download/tools/ReatomeFIs/FIsInGene_070323_with_annotations.txt.zip | reactome.org | 43 |
| SignaLink | v3.1 | http://signalink.org/download | signalink.org | 44 |
| SIGNOR | 3 | https://signor.uniroma2.it/releases/getLatestRelease.php | https://signor.uniroma2.it/ | 45 |
| Spike | 1 | https://www.cs.tau.ac.il/~spike/download/spikeDB.sif.zip | https://www.cs.tau.ac.il/~spike/ | 46 |
| STRING | 12.0 | https://stringdb-static.org/download/protein.links.v12.0/9606.protein.links.v12.0.txt.gz | https://string-db.org/ | 47 |
| TFLink | 1 | https://cdn.netbiol.org/tflink/download_files/TFLink_Homo_sapiens_interactions_All_mitab_v1.0.tsv.gz | https://tflink.net/ | 48 |
| Wan | 1 | http://metazoa.med.utoronto.ca/data/High_confidence_16655_correlations_and_ppi_scores.zip | http://metazoa.med.utoronto.ca/ | 49 |
| Youn | 1 | Supplemental Table 2 | http://dx.doi.org/10.1016/j.molcel.2017.12.020 | 50 |
